# ENDOTHELIAL GENE REGULATORY ELEMENTS ASSOCIATED WITH CARDIOPHARYNGEAL LINEAGE DIFFERENTIATION

**DOI:** 10.1101/2023.10.23.563477

**Authors:** Ilaria Aurigemma, Olga Lanzetta, Andrea Cirino, Sara Allegretti, Gabriella Lania, Rosa Ferrentino, Varsha Poondi Krishnan, Claudia Angelini, Elizabeth Illingworth, Antonio Baldini

**Author notes:** Correspondence to be addressed to Antonio Baldini. email addresses:* IA *OL:*, *AC:* SA *GL:* *RF:* *VPK:* CA *EI:*, AB.

## Abstract

Endothelial cells (EC) differentiate from multiple sources, including the cardiopharyngeal mesoderm, which gives rise also to cardiac and branchiomeric muscles. Here, we used a cardiogenic mesoderm cell differentiation model that also activates an endothelial transcription program to identify endothelial regulatory elements activated in early cardiogenic mesoderm. Integrating our chromatin remodeling and gene expression data with available single-cell RNA-seq data from mouse embryos, we identified 101 putative regulatory elements of EC genes. We then applied a machine-learning strategy, trained on validated enhancers, to predict the probability of the sequences to function as enhancers. The computational assay determined that 50% of these sequences were likely enhancers, some of which have been previously reported. We also identified a smaller set of regulatory elements of well-known EC genes and validated them using genetic and epigenetic perturbation. Finally, we used the integration of multiple data sources and computational tools to search for transcriptional factor binding motifs. In conclusion, we identified novel EC regulatory sequences with a high likelihood to be enhancers, and we validated a subset of them using computational and cell culture models. Motif analyses revealed that the core EC transcription factors GATA/ETS/FOS is a likely driver of EC differentiation in cardiopharyngeal mesoderm.

## BACKGROUND

In mammals, endothelial cells (EC) derive, through vasculogenesis, from the mesoderm but they can differentiate from multiple sources [1,2]. ECs are heterogeneous in their function, transcriptional program, and chromatin landscape; single-cell based studies have provided a wealth of data to this effect [3,4]. Heterogeneity has different causes [5], but, at least in part, it may depend upon lineage of origin as well as epigenomic and enhancer profiles.

The cardiopharyngeal mesoderm (CPM) lineage provides progenitors to various tissues and organs of the lower face, mediastinum and heart [6]. The CPM also provides multipotent progenitors that differentiate into ECs [7–9] [10]. In addition, the second heart field (SHF) [11–13] which derives from the CPM, provides EC progenitors to various components of the cardiovascular system, including ECs of the pharyngeal arch arteries and outflow tract [14,15], which, through endothelial-to-mesenchymal transition, contribute to cardiac valve formation. In particular, *Tbx1* expression, which is a marker of the CPM and SHF, identified these ECs through genetic labeling driven by a *Tbx1^Cre^*allele [15,16]. In addition, time-controlled genetic labeling with an inducible Cre recombinase determined that *Tbx1* was activated in EC progenitors within the time window E7.5-E8.5 in mouse embryos [15]. The molecular events that drive EC differentiation in the CPM are mostly unknown, with some exception [8], and a suitable approach to define them would be to use a dynamic model in which chromatin remodeling and gene expression can be monitored at the critical developmental times. Using such an approach, we found that the differentiation of cardiogenic mesoderm from mouse embryonic stem cells (mESCs) activated an endothelial transcription program. We then measured gene expression and chromatin accessibility in differentiating mESCs within the activation window to identify differentially expressed genes and differentially accessible regions. We then used EC gene information from published single-cell RNA-seq data obtained from *Tbx1^Cre^* and *Mesp1^Cre^* sorted cells from E8.5-E9.5 mouse embryos. Data integration and analysis identified and computationally scored 101 putative regulatory elements activated in the *Tbx1^Cre^*-selected EC cluster. Finally, we identified and validated putative regulatory elements associated with a small set of well-known EC genes.

In summary, our results provide a novel and systematic experimental approach to identify cell type-specific regulatory elements during differentiation, and the results obtained shed light onto the EC regulatory elements activated during cardiogenic mesoderm differentiation.

## RESULTS

### Cardiac mesoderm differentiation of mouse embryonic stem cells activates endothelial differentiation and it can be driven to yield a nearly homogeneous EC population

We have used a published protocol to derive cardiac mesoderm from mouse embryonic stem cells (mESCs) [17] (Fig. 1A). We found expression of endothelial genes at differentiation day 4 (d4), while these genes were not detected at d2 (Fig. 1B). RNA-seq analysis performed on two replicates from two independent differentiation experiments confirmed the activation of an endothelial expression program within this time window (Fig. 1C, Tab. 1, Additional file 1: Tab. S1). Therefore, we used the d2-d4 time window in our search for EC enhancers.

**Figure 1.**
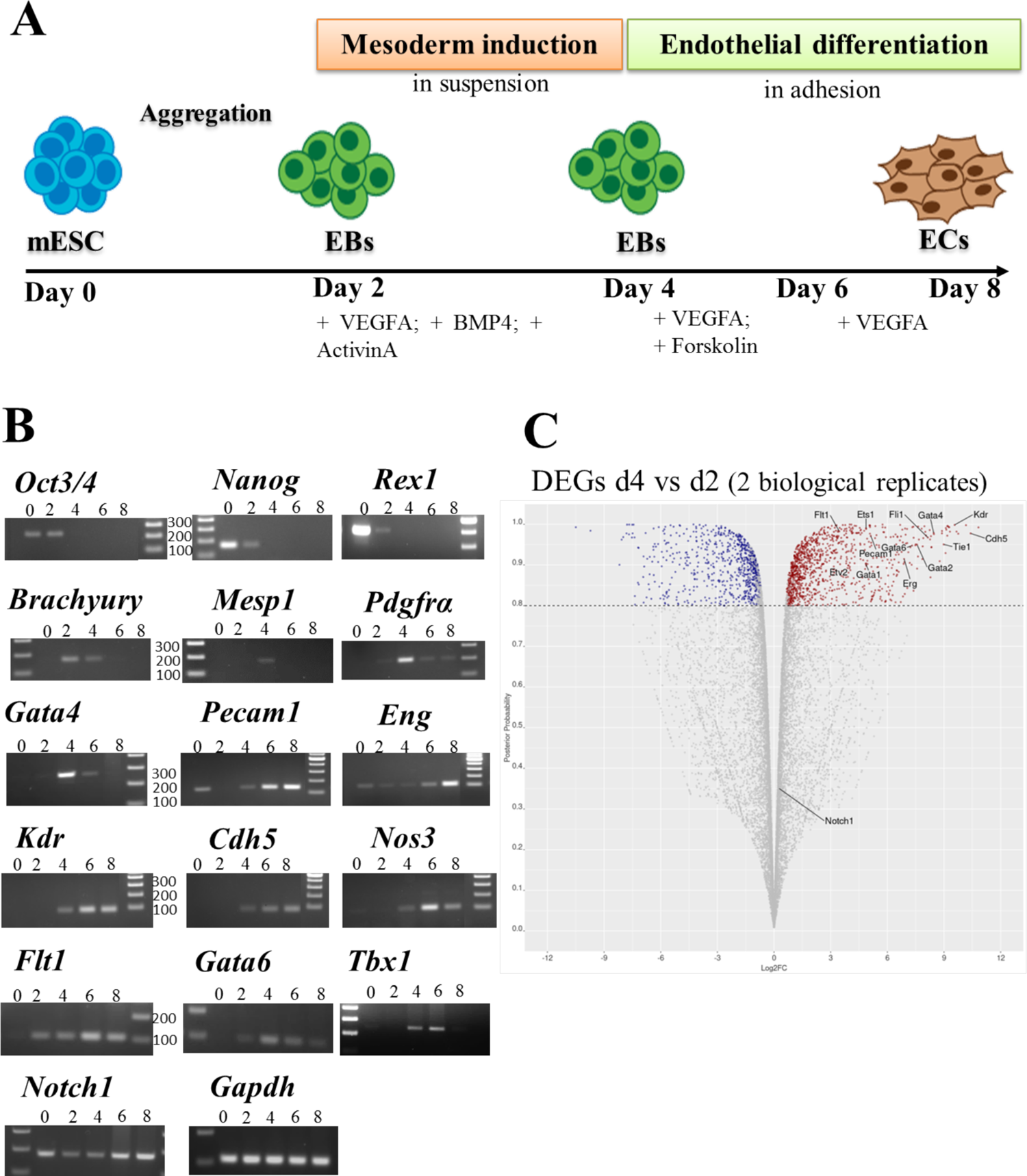
Activation of a EC transcription program during cardiogenic mesoderm differentiation of mESC. **(A)** Schematic illustration of the EC differentiation protocol from mESCs. **(B)** Expression of pluripotency (*Oct3/4; Nanog; Rex1*), mesodermal (*Brachyury; Mesp1; Pdgfrα; Gata4*), endothelial marker genes (*Pecam1; Eng; Kdr; Cdh5; Nos3; Flt1; Gata6; Notch1*), and the CPM marker *Tbx1* during differentiation by RT-PCR. *Gapdh* was used as normalizer. The molecular weight marker is the 100bp ladder. **(C)** RNA-seq volcano plot of differentially expressed genes (DEGs) in d4 vs d2 samples in two biological replicates. Genes downregulated at d4 (n. 731) are indicated in blue, genes up regulated are in red (n. 1088). We indicate examples of endothelial marker genes (*Flt1; Kdr; Ets1; Fli1; Gata4; Pecam1; Cdh5; Etv2; Gata6; Gata1; Erg; Tie1; Gata2, Notch1*).

**Table 1.** Gene ontology of genes significantly up regulated at d4 compared to d2.

| source | term | term id | adj p value | term size | query size | intersection |
| --- | --- | --- | --- | --- | --- | --- |
| **GO:B**<br>**P** | anatomical structure morphogenesis | GO:0009653 | 1.84E-52 | 1999 | 1082 | 353 |
| **GO:B**<br>**P** | circulatory system development | GO:0072359 | 8.39E-52 | 900 | 1082 | 217 |
| **GO:B**<br>**P** | anatomical structure development | GO:0048856 | 5.11E-50 | 4047 | 1082 | 549 |
| **GO:B**<br>**P** | multicellular organismal process | GO:0032501 | 1.56E-49 | 4609 | 1082 | 597 |
| **GO:B**<br>**P** | anatomical structure formation involved in morphogenesis | GO:0048646 | 1.75E-48 | 877 | 1082 | 208 |
| **GO:B**<br>**P** | **vasculature development** | GO:0001944 | 7.91E-48 | 583 | 1082 | 164 |
| **GO:B**<br>**P** | cell surface receptor signaling pathway | GO:0007166 | 1.37E-47 | 1786 | 1082 | 318 |
| **GO:B**<br>**P** | **blood vessel development** | GO:0001568 | 2.07E-47 | 556 | 1082 | 159 |
| **GO:B**<br>**P** | developmental process | GO:0032502 | 2.07E-47 | 4400 | 1082 | 573 |
| **GO:B**<br>**P** | signal transduction | GO:0007165 | 1.35E-45 | 3507 | 1082 | 488 |

Flow cytometry using the endothelial-specific marker VE-Cadherin, encoded by the *Cdh5* gene, revealed that at d2 there were no detectable VE-Cadherin+ cells, while at d4, a small percentage (19 %) was present (Fig. 2A). Therefore, we extended the differentiation protocol in order to increase the EC population. To this end, at d4 we added a high concentration of VEGFA (200ng/ml) and Forskolin (2µM), as suggested by a published protocol [18]. Following this treatment, at d6 and d8, the percentage of VE-Cadherin+ cells increased to 57.4% and 91.3%, respectively (Fig. 2A). Similar results were obtained in multiple experiments. Next, we performed Matrigel assays on d8 cells [19] in order to determine whether they formed the tubule-like networks expected of fully differentiated ECs. Results indicated that d8 cells had this capacity (Fig. 2B). Thus, this modified differentiation protocol produced a nearly homogeneous population of ECs.

**Figure 2.**
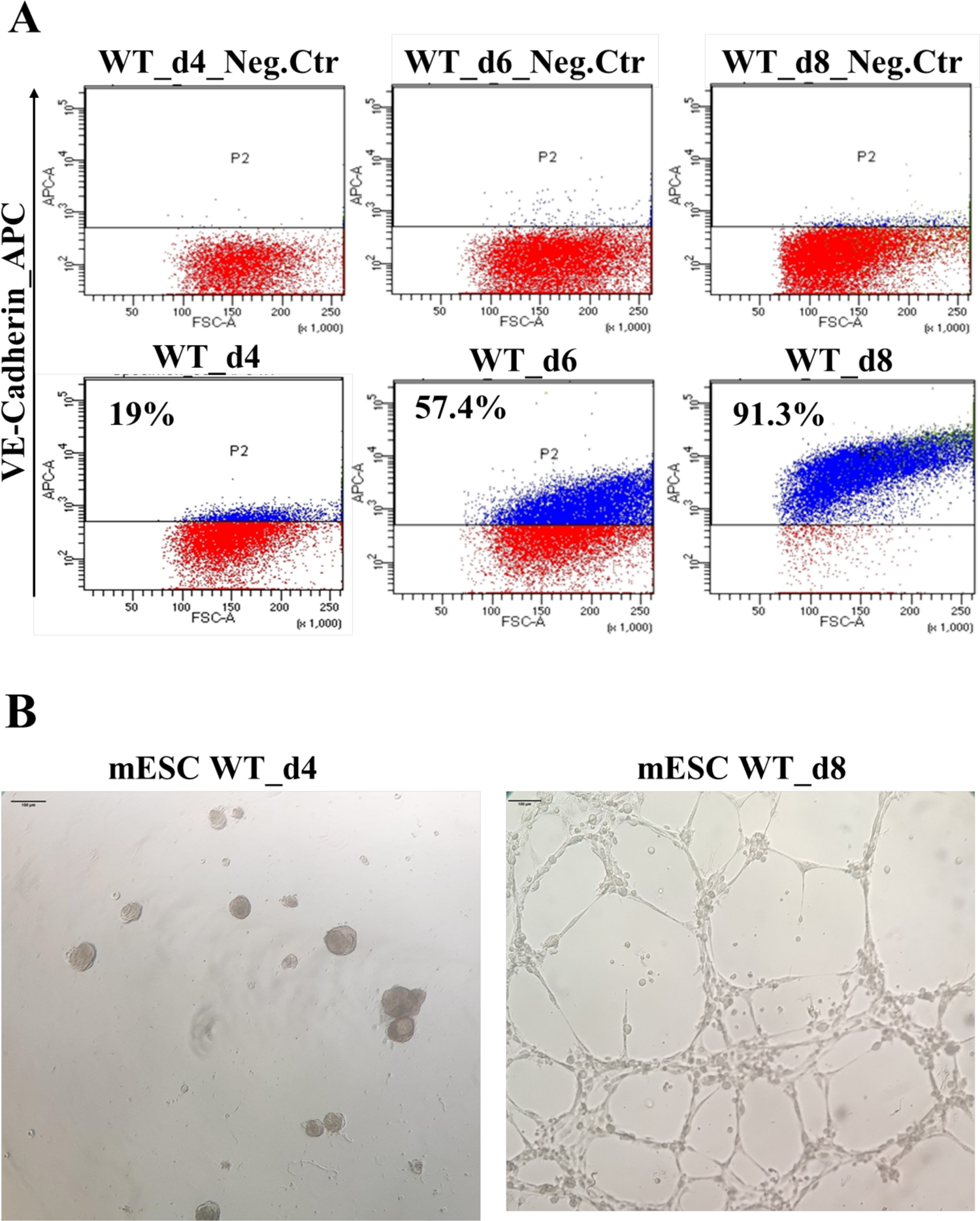
Progressive EC differentiation from cardiogenic mesoderm. **(A)** Flow cytometry using anti-VE-Cadherin antibody during mESC differentiation. The VE-Cadherin^+^ subpopulation is identified at day4-6-8 of differentiation. Negative control is isotype IgG1 control antibody-labeled differentiating cells. **(B)** *In vitro* tube formation assay (Matrigel) of d4 cells (left, negative control) and d8 cells (right) plated for 24h. The scale bar is 100µm.

### An unbiased strategy identifies putative regulatory elements in early EC differentiation

Having identified the d2-d4 time window for the activation of an EC transcription program, we performed ATAC-seq at these two time-points in order to localize regions of dynamic chromatin accessibility across the genome. Experiments were performed in two biological replicates (from two independent differentiation experiments) and we subsequently considered consensus peaks only, i.e., peaks that were called in both replicates. Thus, we identified a total of 20268 consensus peaks at d2 and 17110 at d4 (Additional file 2: Tab. S2), of which 8773 were differentially accessible regions (DARs) as determined by the DiffBind and Descan2 software tools [39; 40]; again we only considered DARs that were identified by both tools (Figs. 3A and 3B). We then derived a list of marker genes of an EC cluster obtained from single-cell RNA-seq (scRNAseq) experiments performed on cells selected using a *Tbx1Cre* driver combined with a GFP reporter. *Tbx1* is a CPM marker and cells were FACS-purified from E8.5 and E9.5 mouse embryos [20]. The EC cluster was characterized by 252 marker genes (listed in Additional file 3: Tab. S3) that we intersected with our dataset of d2-d4 DARs opened at d4 (n=4408) (Fig. 3A). This resulted in 101 regions that were significantly more accessible at d4, compared to d2, and were associated with EC expressed genes (Tab. 2). We then performed a computational prediction of the probability that these regions are enhancers. To this end, we used a machine-learning procedure based upon a logistic regression test trained on validated enhancer sequences (see methods for details). The average scores obtained with this procedure are reported in Tab. 2. Of the 101 regions identified, 57 scored more than 0.5, indicating significant likelihood of being enhancers; of these, 15 (26%) have been reported in the literature (references indicated on Tab. 2). A search for transcription factor binding motifs within the 101 regions identified significantly enriched motifs (as the background, we used the peakome associated with expressed genes). Specifically, we found motifs of GATA, ETS transcription factor families, and FOS, a subunit of the AP-1 transcription complex (Fig. 3C and Additional file 4: Tab. S4). GATA and ETS factors are co-present in 55% (56 out of 101) of the regions tested. The expression of *Gata1*, *Gata2*, *Gata4*, *Gata6, Fos and Erg*, as well as other ETS family members (e.g. *Ets1, Ets2, Etv2, Fli1, Elk1, Elf1*) genes were strongly up regulated at d4, relative to d2 (Additional file 1: Tab. S1).

**Figure 3.**
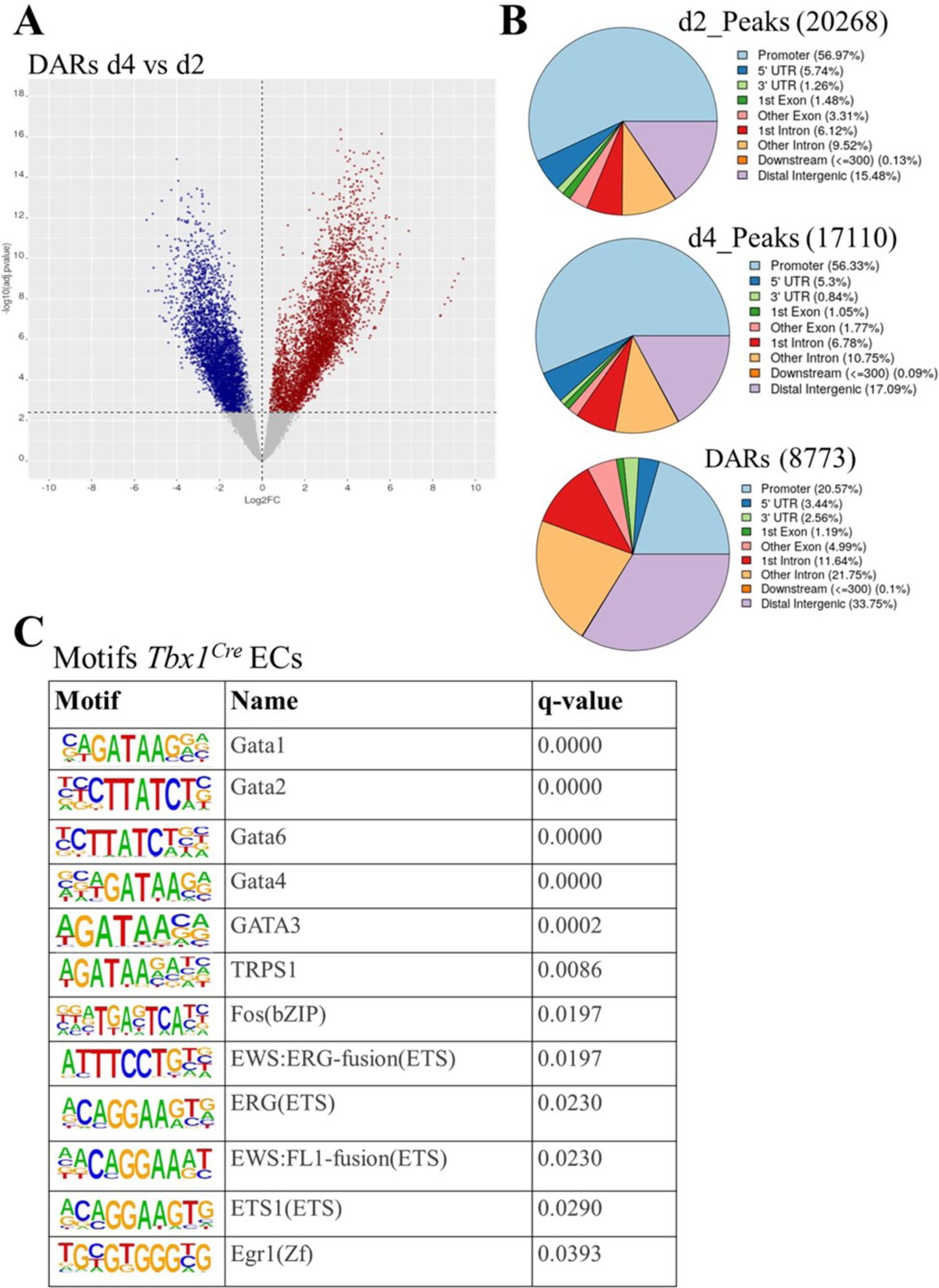
Chromatin remodeling during mESC differentiation. **(A)** Volcano plot of differentially accessible regions (DARs) in d4 vs d2 samples. In blue are DARs decreased at day4; in red are DARs increased at day4. **(B)** Distribution of total ATAC peaks at d2, d4, and DARs relative to gene features. The promoter region has been set at ±1000bp to transcription start site (TSS). **(C)** Enriched known motifs evaluated by HOMER using DARs mapped to marker genes of the EC cluster reported by Nomaru et al., 2021 selected from the *Tbx1^Cre^*-sorted population of mouse embryos at E8.5 and E9.5. The full motif search results are reported on Additional file 4: Tab. S4.

**Table 2.**
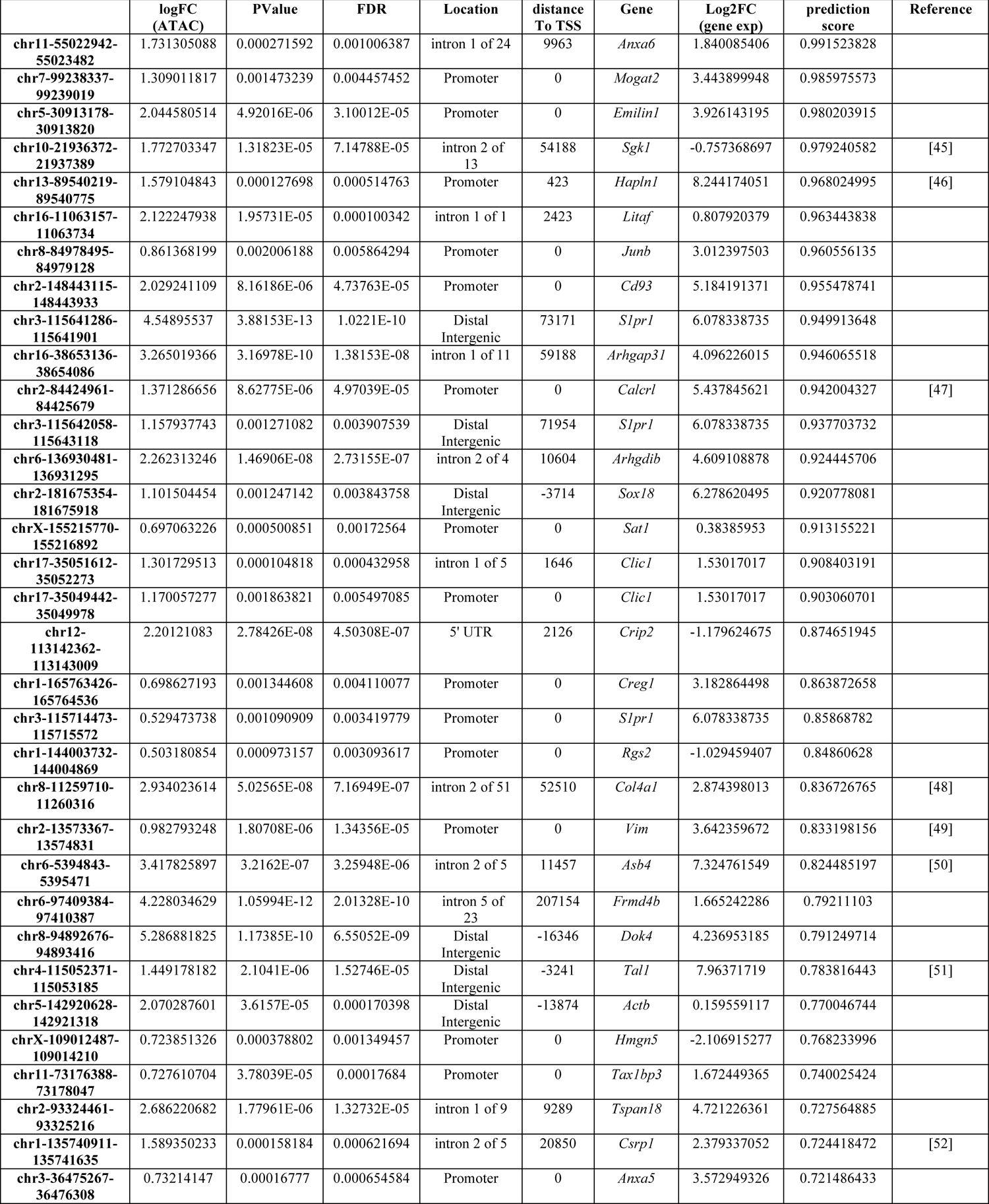
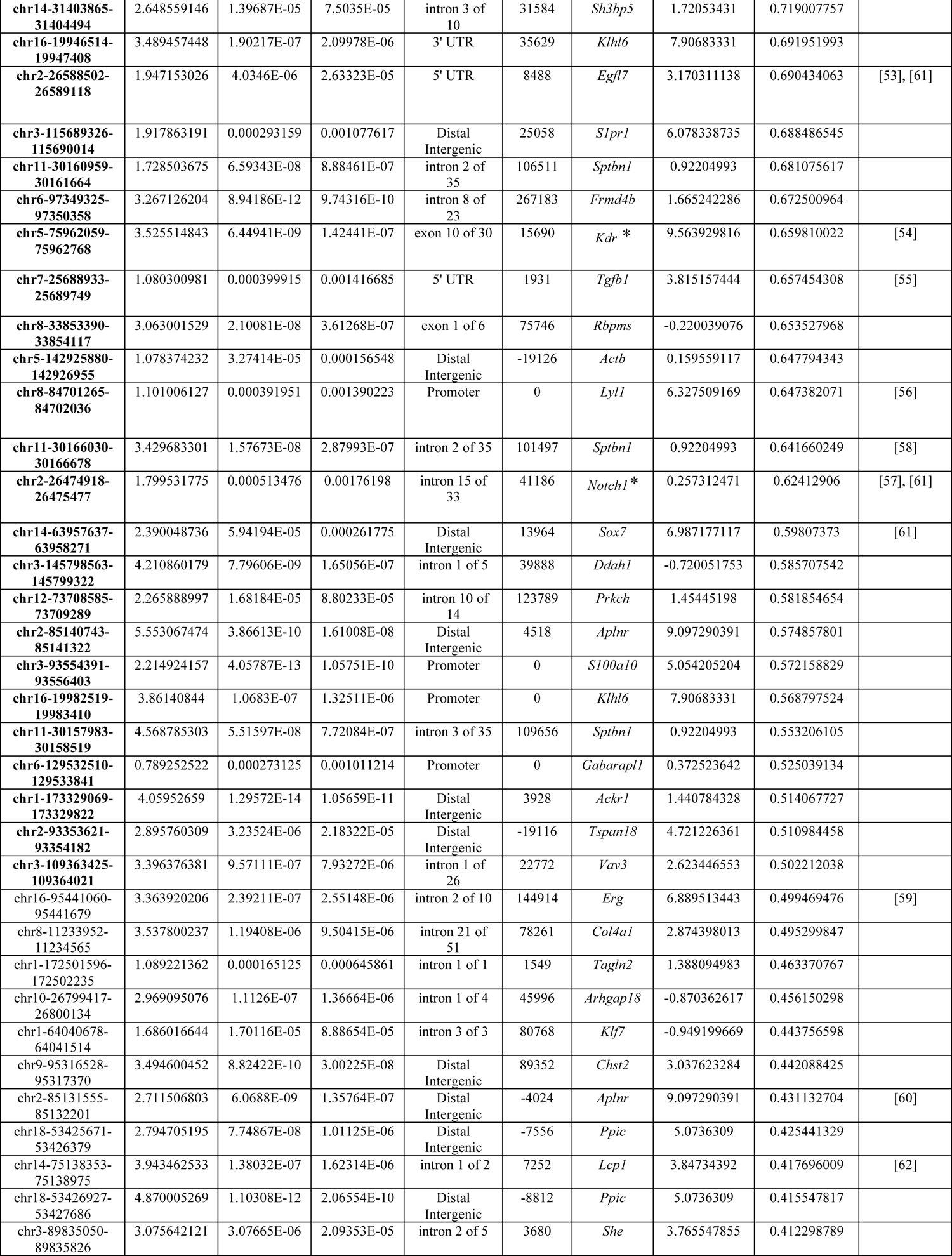
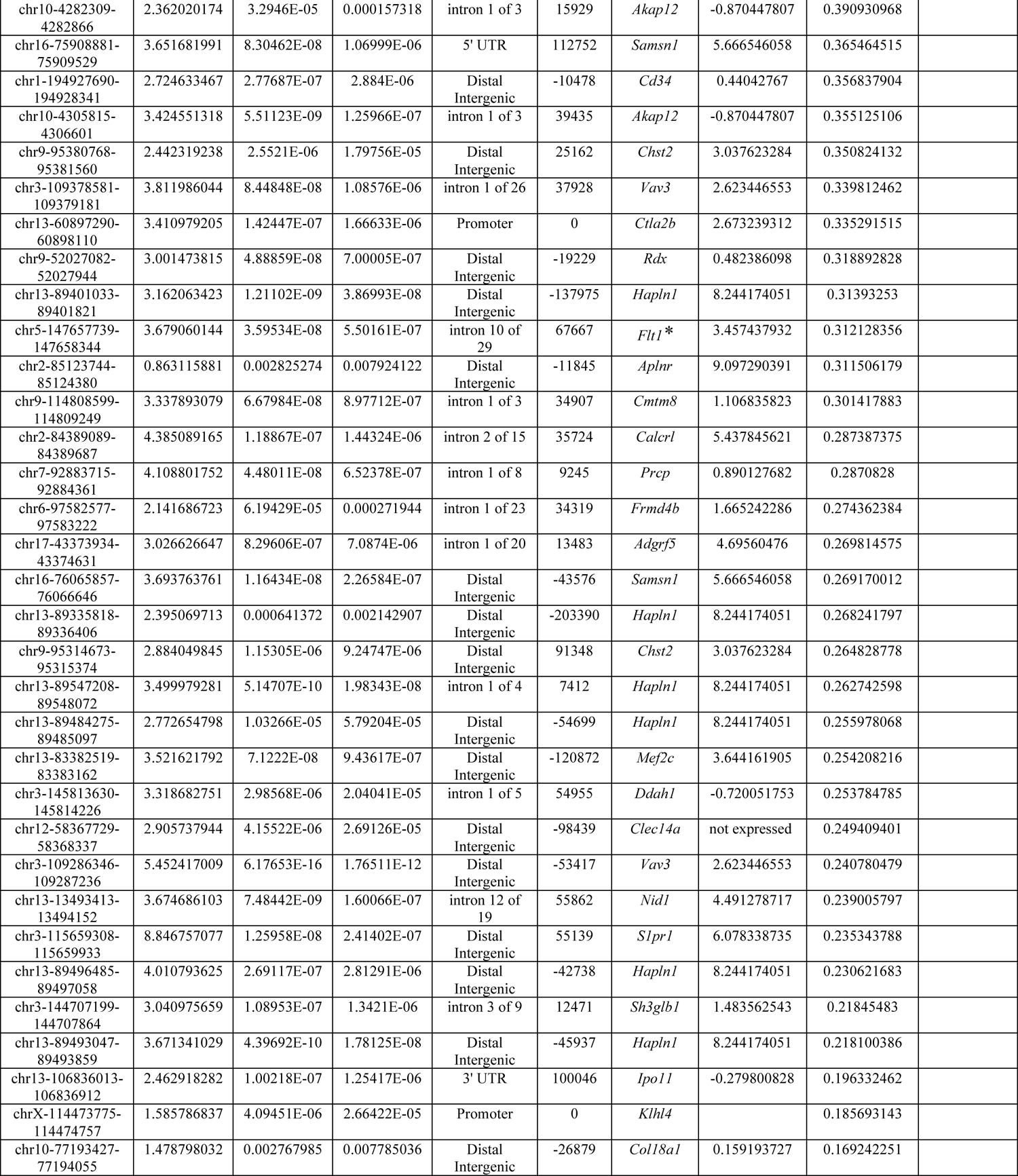
Chromatin regions opened at d4 and mapped to EC marker genes of a *Tbx1^Cre^* - selected EC cluster.

Next, we repeated the procedure using marker genes expressed in endothelial clusters of a scRNA-seq dataset from *Mesp1^Cre^*-sorted cells at the same developmental stage (E9.5) [20]. *Mesp1^Cre^*-sorted cells include ECs derived from the entire anterior mesoderm and not just the cardiopharyngeal mesoderm, practically the entire vascular bed of the trunk, as the head and most posterior regions of the embryos were removed before sorting [20]. In this study, two endothelial clusters were identified, named c2 and c16, that shared 801 marker genes (Additional file 3: Tab. S3). We mapped DARs upregulated at d4 to these sets of genes and identified 536 unique putative regulatory elements, of which, 283 (52.8%) had a score above 0.5 (Additional file 3: Tab. S3). We conducted motif searches using DARs upregulated at d4 mapped to marker genes of the two endothelial clusters separately, 434 regions for *Mesp1^Cre^* c16 and 367 regions for *Mesp1^Cre^* c2. Results identified a more extensive set of motifs than the one identified using the *Tbx1^Cre^* dataset, but the most enriched ones were again GATA and ETS factors (Additional file 4: Tab. S4).

### Identification and validation of EC regulatory elements (RE) associated with major EC differentiation genes

Next, we applied a different approach to the identification of EC-REs: we selected a group of well-known endothelial genes that were expressed at d4, on the basis of RNA-seq data, and that exhibited regions of increased chromatin accessibility at d4. We focused on 6 putative, non-promoter REs associated with six genes: *Kdr* (encoding VEGFR2), *Chd5* (encoding VE-CADHERIN), *Eng* (encoding ENDOGLIN), *Flt1* (encoding VEGFR1), *Pecam1*, and *Notch1*. Computational prediction indicated that four of the six putative EC-REs identified had a score above 0.5, indicating a high probability to be enhancers (Tab. 3, Fig. 4). Furthermore, the two putative EC-REs associated with *Kdr* and *Notch1* are amongst the 101 regions open at d4 and mapped to EC marker genes (asterisks on Tab. 2).

**Figure 4.**
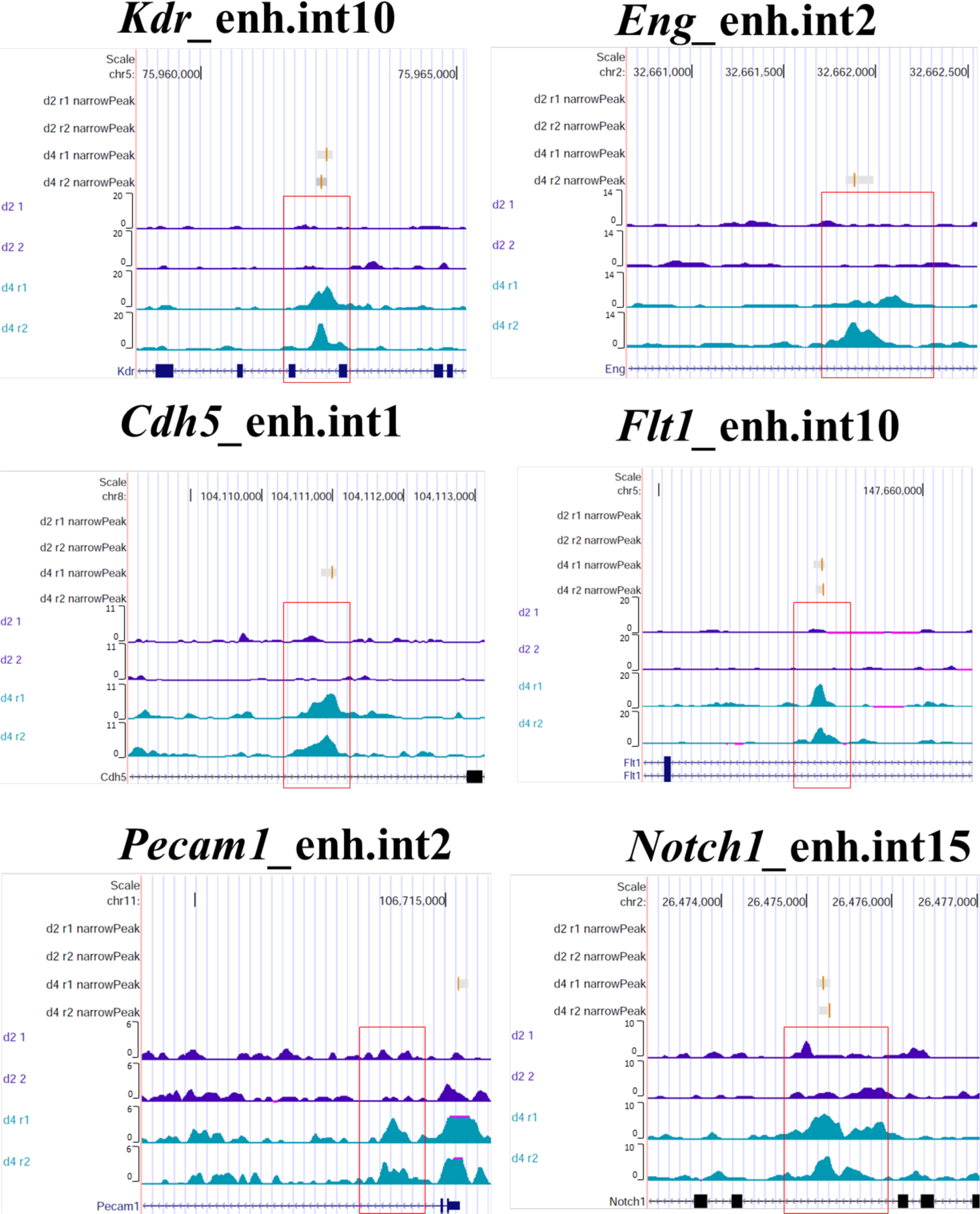
Selection of putative regulatory elements in major EC genes (EC-REs). ATAC-seq coverage associated to 6 putative EC-REs, associated to 6 selected EC genes: *Kdr; Eng; Cdh5; Flt1; Pecam1; Notch1.* On vertical axis there are the genome coverage of d2 (replicate1 and replicate2) and d4 (replicate1 and replicate2). Red boxes indicate the open chromatin region at d4.

**Table 3.**
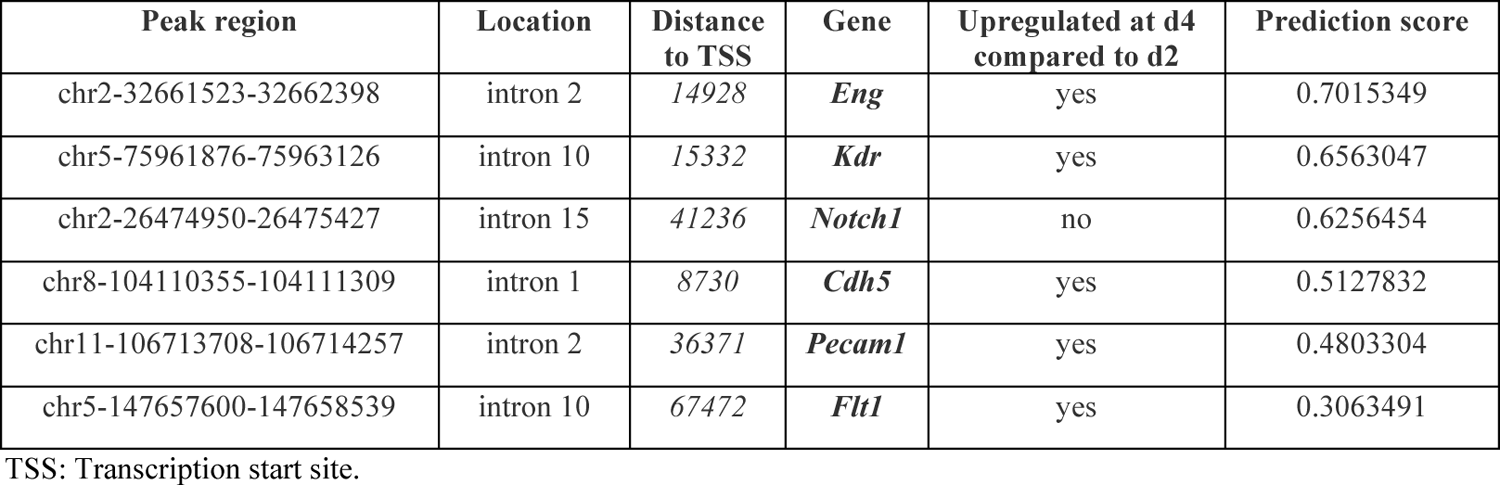
Putative EC regulatory regions associated with the genes indicated.

To test the importance of the putative EC-REs, we used an epigenetic reprogramming strategy based on CRISPR-dCAS9:LSD1 (Fig. 5A). We first generated an ES cell line that stably expressed the dCAS9:LSD1 construct (named #B1 dCas9-LSD1). We then designed crRNAs targeting the six EC-REs and a control crRNA targeting a gene desert sequence (Additional file 5: Tab. S5). We transfected #B1 dCas9-LSD1 cells with targeting and control gRNAs complex (crRNAs:ATTO-tagged tracrRNA), FACS-purified the ATTO+ cells, and subjected them to the EC differentiation protocol. Cells were harvested at d4, d6, and d8 and the expression of the targeted genes was measured by quantitative real time PCR (qPCR). Experiments were repeated at least four times. Results showed that LSD1 targeting of the six EC-REs resulted in reduced expression of the associated genes (Fig. 5B), with the exception of the *Pecam1*-associated RE, which also had a low probability score (Tab. 3). In most cases, the reduction in gene expression was more evident at the later stages of differentiation tested, namely d6 and, even more so at d8.

**Figure 5.**
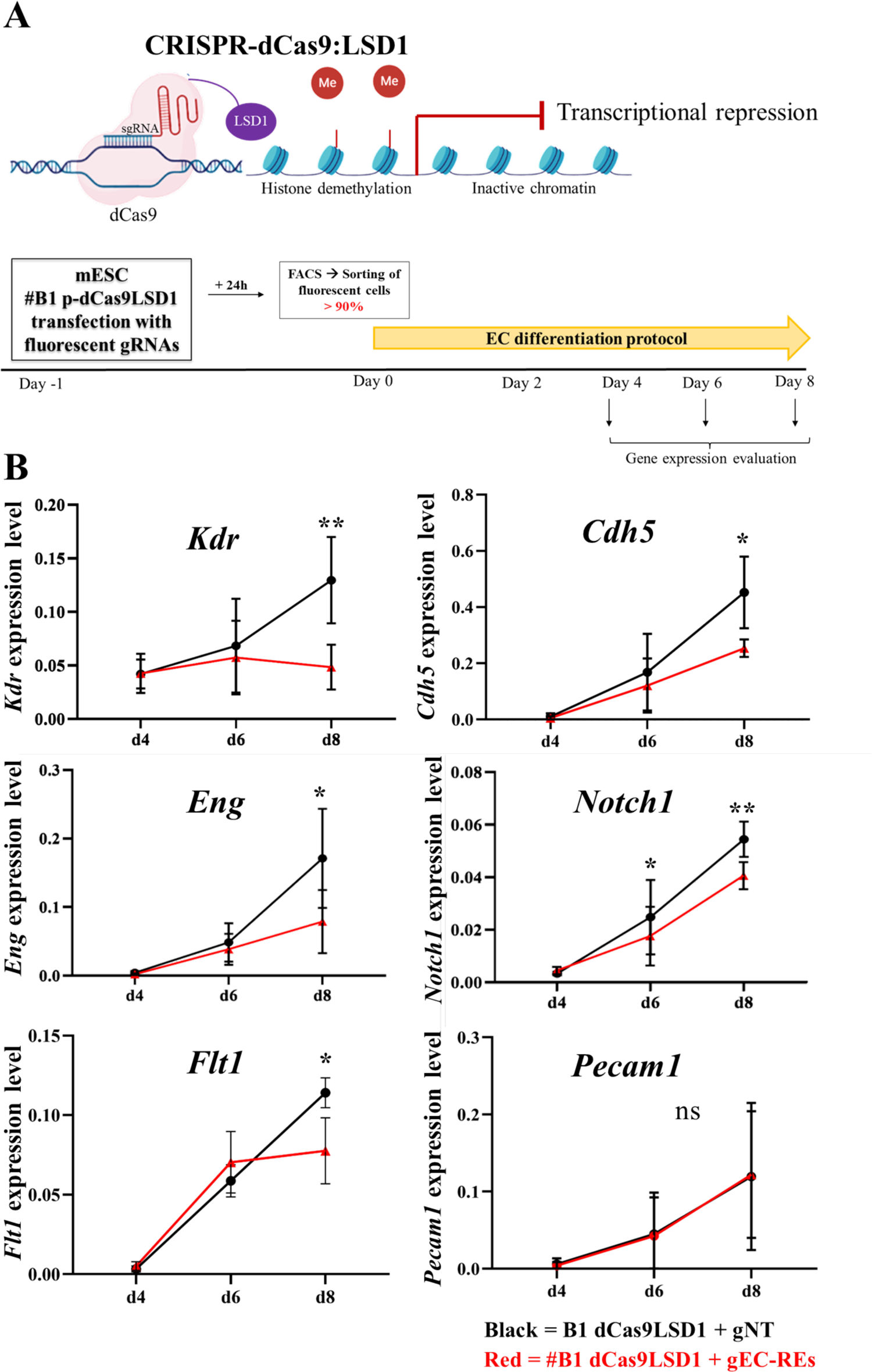
Epigenetic reprogramming validates putative regulatory elements. **(A)** *Top*: Schematic overview of CRISPR-dCas9:LSD1 system: the fusion protein dCas9:LSD1 is able to bind DNA target and LSD1 can demethylate histone H3 lysine 4 (H3K4me1 and me2) near the putative enhancer region to decommission the enhancer. *Bottom*: cartoon of the experimental plan. mESC #B1 dCas9LSD1 were transfected with fluorescent gRNAs. Fluorescent sorted cells were differentiated into ECs from day0 to day8. Samples were collected at day4, day6 and day8 to analyze the gene expression. **(B)** Quantitative real time PCR (qPCR) analysis of *Kdr; Cdh5; Eng; Notch1; Flt1* and *Pecam1* mRNA expression level in cells of clone #B1 dCas9LSD1 transfected with gRNAs targeted (in red) or control (in black) during EC differentiation. X-axis denotes the three time points (d4-d6-d8); y-axis indicates the expression level, evaluated using the 2^−ΔCt^ method. *Gapdh* expression was used as normalizer. Values are the average of four (n=4) biological replicates ± standard deviation (SD). p-value (*) < 0.05 and p-value (**) < 0.01 are considered significant; ns= no statistical significance (parametric paired t-test, one-tailed).

### The EC-REs for Notch1 and Pecam1 are required for gene expression

Next, we selected the *Notch1* and *Pecam1* EC-REs for further testing based on the importance of the associated genes for EC differentiation, and because of the negative results obtained with epigenetic reprogramming for *Pecam1*. We deleted the putative EC-REs using CRISPR-Cas9; for each RE we selected two gRNAs flanking the segment (Fig. 6A, gRNA sequences are shown in Additional file 5: Tab. S5), and these were transfected into mESCs along with the Cas9 protein and ATTO-labeled tracrRNA. We then plated FACS-purified ATTO+ cells to a clonal density. Clones were later picked and expanded into 96-well plates. DNA extracted from clones was screened by PCR to identify clones carrying homozygous deletion of the putative RE. We expanded two homozygously deleted clones for each deleted RE. All four clones were used in multiple differentiation experiments (n=5). Results showed that both *Pecam1* and *Notch1* expression were significantly affected by the deletion of their respective REs at d6 and at d8 (Fig. 6B).

**Figure 6.**
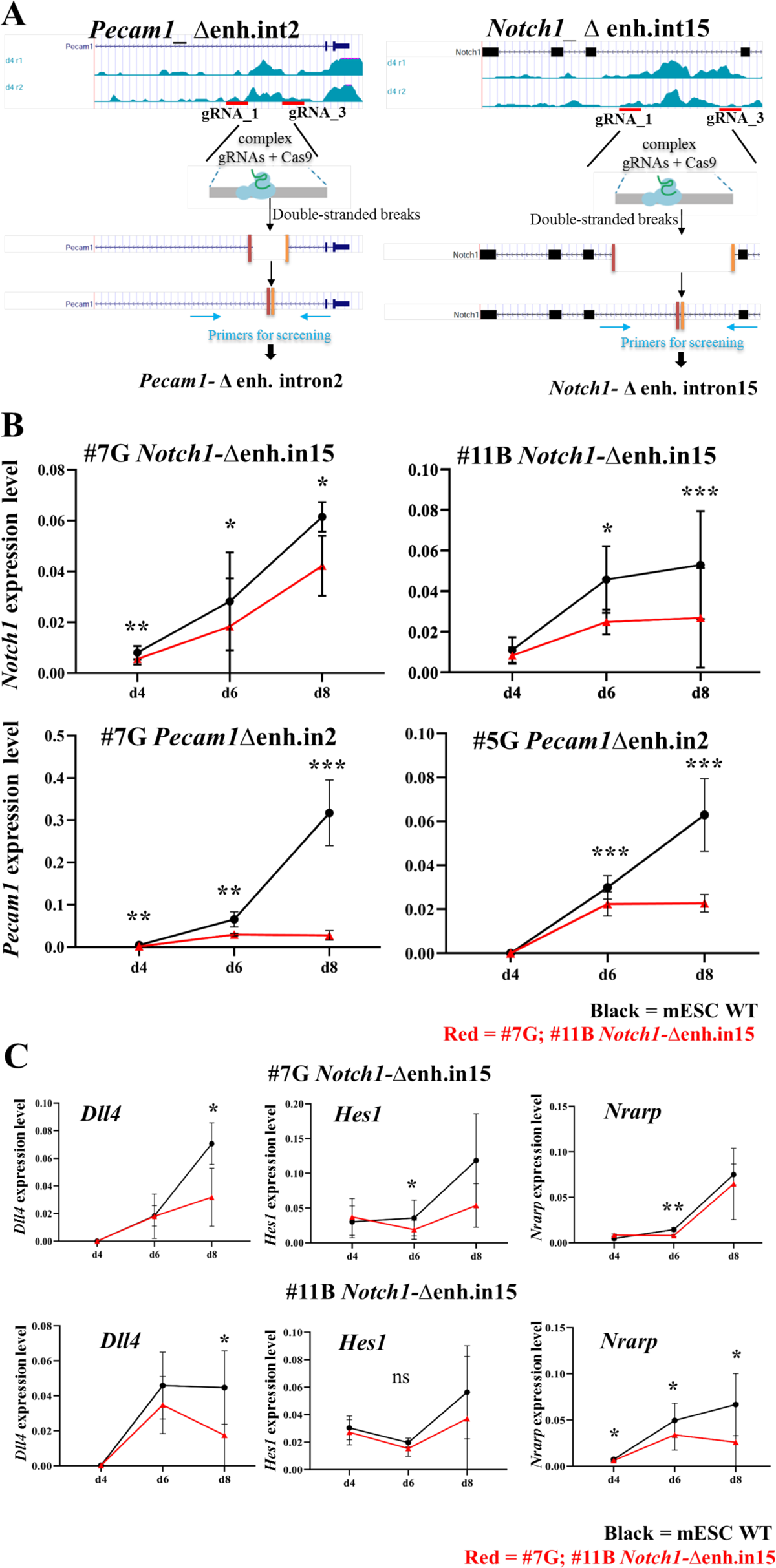
Homozygous deletion of putative regulatory elements of Notch1 and Pecam1 genes reduced their expression during EC differentiation. **(A)** Scheme of the steps of targeted Pecam1-enh.int2 and Notch1-enh.int15 deletion with CRISPR/Cas9. Red lines indicate the position of the two gRNAs used. **(B)** Quantitative real time PCR (qPCR) analysis of Notch1 mRNA expression level in mESC Notch1-Δenh.in15 (clones #7G; #11B) and Pecam1 in mESC Pecam1Δenh.in2 (clones #7G; 5G) during EC differentiation. Notch1 and Pecam1 expression was reduced in mutant cell lines (in red), compared to WT cells (in black), used as control. X-axis denotes the three time points (d4-d6-d8); y-axis indicates the expression level, evaluated using the 2^−ΔCt^ method. Gapdh expression is used as normalizer. Values are the average of five biological replicates ± standard deviation (SD). p-value (*) < 0.05; p-value (**) < 0.01 and p-value (***) < 0.001 are considered significant; ns= no statistical significance (parametric paired t-test, one-tailed). **(C)** Quantitative real time PCR (qPCR) analysis of gene expression of Notch1-related genes in mESC Notch1-Δenh.in15 (clones #7G; #11B) during EC differentiation. p-value (*) < 0.05 and p-value (**) < 0.01 are considered significant; ns= no statistical significance (parametric paired t-test, one-tailed).

We next tested whether the deletion of the *Notch1* EC-RE affected expression of a subset of NOTCH1 target genes, namely *Hes1, Nrarp*, and *Dll4*. Results showed that all of these genes were affected by the deletion of the enhancer, but with some differences. Specifically, *Hes1* and *Nrarp* were significantly and consistently down regulated at d6 but not at d8. Conversely, *Dll4* was down regulated only at d8 (Fig. 6C). Thus, deletion of the EC-RE identified here was sufficient to cause dysregulation of at least part of the NOTCH1 signaling pathway.

## DISCUSSION

The cardiopharyngeal mesoderm [6] is a lineage that provides progenitors to various structures including those of the heart, pharyngeal apparatus, and vessels [9,10]. Endothelial cells are heterogeneous in origin and single cell sequencing assays are starting to define specific transcription and chromatin profiles depending on the tissue of origin [4]. However, whether cells destined to differentiate in EC are primed by distinct mechanisms according to their origin, is still unclear. One possible avenue to address this question is to identify tissue-specific enhancers for each lineage. In this study, we propose an approach that leverages novel and published data, integrated with software tools, and genetic/epigenetic editing in a cell differentiation model. This integrated approach identified a group of putative EC enhancers, some of which had already been reported in the literature and were validated in our model. For this study, we used a mesoderm differentiation protocol originally proposed for cardiogenic mesoderm induction [17] and observed the activation of EC-specific gene expression and chromatin remodeling (as assayed by ATAC-seq) 48hrs after induction. However, at this stage (d4) only a small percentage of cells exhibit an EC phenotype as defined by the expression of VE-cadherin on the cell surface, suggesting that activation of the EC program is at an early stage. Boosting VEGF signaling after mesoderm induction promoted EC differentiation such that a near-homogeneous EC-like population was obtained at d8, as measured by VE-cadherin expression and Matrigel assays. We selected to leverage chromatin dynamics of EC-specific gene activation at an early developmental time window (d2-d4) in order to capture the regulatory sequences associated with the activation of the EC program in cardiogenic mesoderm. To this end, we used data-rich bulk ATAC-seq and RNA-seq information, combined with published high-resolution tissue- and time-specific scRNA-seq of cells that were selected using the *Tbx1^cre^* driver, *Tbx1* being a marker of the cardiopharyngeal mesoderm. This enabled us to identify 101 regions that became more accessible in the selected time window and mapped to EC genes defined by scRNA-seq data (Tab. 2). Of the 57 putative enhancers scoring > 0.5 identified through our unbiased approach, 15 (26%) were already reported in the literature, thus suggesting that our approach was efficient in detecting likely regulatory elements in our model. In addition, motif analysis showed enrichment of transcription factors known to be involved in EC differentiation, further supporting the suitability of the approach. However, we did not validate these putative enhancer directly in our model, thus further work will be necessary to establish the reliability of our approach for systematic identification of cell type-specific enhancer sequences.

The candidate gene approach was designed to identify regulatory sequences “activated” in our model and associated with genes known to be involved in EC differentiation. We have validated them regardless of the prediction score and found five of the six tested putative REs to regulate the respective genes. Overall, we identified regulatory elements for many of the known genes involved in EC differentiation, including a subset of genes expressed in CPM-derived cells in vivo like for example *Notch1* [8].

Epigenetic reprogramming, while providing consistent results, proved to be variable in our hands. Sources of variability may be the efficiency of transfection, the gRNAs, or perhaps variable extent of chromatin modification induced by the dCAS9:LSD1 complex. Furthermore, the inconsistent results obtained with the *Pecam1* putative enhancer using epigenetic reprogramming and gene editing may be due to different reasons. We speculate that perhaps the sequence is not a regulatory element (as suggested by the low prediction score) but the genetic deletion may have altered the expression of the gene by interfering with processes like RNA maturation/splicing or other structural perturbation of the gene.

The dCas9-recruited repressor could potentially cause chromatin modifications beyond the intended targeted sequence, particularly if the promoter is nearby. The six enhancers tested with this method are all fairly distant from the transcriptions start site (TSS, Tab. 3). The closest is 8.7Kb from the TSS, the others are between 15 and 67 Kb.

Searches for consensus sequences in the putative enhancer regions identified GATA motifs as the most enriched, together with ETS factors (ERG, FLI,) and the AP-1 subunit FOS (Fig. 3B). EC genes from cell clusters derived using the *Mesp1^Cre^* driver [20], which captures a larger and more diverse EC population than *Tbx1^Cre^*, exhibited DARs enriched for a more extensive set of motifs, but they also included GATA and ETS motifs. Interestingly, the *Mesp1^cre^*dataset motifs also included transcription factor families that play a role in CPM development, such as T-BOX, FOX, and MEIS factors (Additional file 4: Tab. S4), raising the question of whether they may be involved in enabling the EC transcription program in the CPM.

Overall, the results of consensus sequence searches suggest that there is a core of transcription factors, GATA-ETS-FOS, that are central to the activation of the EC program in our model. There is ample literature indicating GATA and ETS transcription factors as general players in endothelial differentiation (reviewed in De Val and Black, 2009). It is therefore possible that during mesoderm induction in our system, GATA factors act as pioneers to establish the conditions necessary for the binding of other, lineage determining factors, such as ETS/ERG. This is consistent with the established role of GATA factors as pioneer transcription factor (review in [21,22]) and the role of ETS factors as core transcription factors in endothelial differentiation [3,23–27].

## Conclusions

Overall, our strategy efficiently identified putative enhancers of cell-type specific genes during differentiation. It provided us with an extensive list of regulatory sequences with probability scores calculated using a machine-learning approach. Furthermore, it allowed us to identify and validate a smaller set of regulatory sequences of well-known genes involved in EC differentiation. The identification and validation strategies applied here are applicable to other cell types, whenever a suitable differentiation model is available, although more extensive bench validation experiments will be required before the proposed approach may be considered an established pipeline for enhancer identification.

## MATERIALS AND METHODS

### Mouse embryonic stem cells (mESC) culture and manipulation

ES-E14TG2a mESCs (ATCC CRL-1821) were cultured without feeders and maintained undifferentiated on gelatin-coated dishes in GMEM (Sigma Cat# G5154) supplemented with 10^3^ U/ml ESGRO LIF (Millipore, Cat# ESG1107), 15% foetal bovine serum (ES Screened Fetal Bovine Serum, US Euroclone Cat# CHA30070L), 0.1 mM non-essential amino acids (Gibco, Cat# 11140-035), 0.1 mM 2-mercaptoethanol (Gibco, Cat# 31350-010), 0.1 mM L-glutamine (Gibco, Cat# 25030081), 0.1 mM Penicillin/Streptomycin (Gibco, Cat# 10378016), and 0.1 mM sodium pyruvate (Gibco, Cat# 11360-070). Cells were passaged every 2–3 days using 0.25% Trypsin-EDTA (1X) (Gibco, Cat# 25200056) as the dissociation buffer. For differentiation, E14-Tg2a mESCs were dissociated with Trypsin-EDTA and cultured at 75,000 cells/ml in serum-free media: 75% Iscove’s modified Dulbecco’s media (Cellgro Cat# 15-016-CV) and 25% HAM F12 media (Cellgro #10-080-CV), supplemented with N2 (GIBCO #17502048) and B27 (GIBCO #12587010) supplements, penicillin/streptomycin (GIBCO #10378016), 0.05% BSA (Invitrogen Cat#. P2489), L-glutamine (GIBCO #25030081), 5 mg/ml ascorbic acid (Sigma A4544) and 4.5 × 10^-4^M monothioglycerol (Sigma M-6145). After 48h in culture, the EBs were dissociated using the Embryoid Body dissociation kit (cod. 130-096-348 Miltenyi Biotec) according to the manufacturer’s protocol and reaggregated for 40h in serum-free differentiation media with the addition of 8 ng/ml human Activin A (R&D Systems Cat#. 338-AC), 0.5 ng/ml human BMP4 (R&D Systems Cat# 314-BP), and 5 ng/ml human VEGF (R&D Systems Cat#. 293-VE). The 2-day-old EBs were dissociated and 6 × 10^4^ cells were seeded onto individual wells of a 24-well plate coated with 0.1% gelatine in EC Induction Medium consisting of StemPro-34 medium (Gibco #10639011), supplemented with SP34 supplement, L-glutamine, penicillin/streptomycin, 200 ng/ml human-VEGF, and 2 μM Forskolin (Abcam, ab120058). The Induction Medium was changed after one day. On day six of differentiation, the cells were dissociated and replated on 0.1% gelatine-coated dishes at a density of 25,000 cells/cm^2^ in EC Expansion Medium, consisting of StemPro-34 supplemented with 50 ng/ml human-VEGF. Stem cell derived endothelial cells were maintained until they reached confluency (about 2-3 days). EC Expansion Medium was replaced every other day.

For the Matrigel assay, 400µL of Matrigel (BD Matrigel Basement Membrane Matrix Growth Factor Reduced, Phenol Red Free cat. 356231) was aliquoted into each well of a 12-well plate and incubated for 30–60 min at 37 °C to allow the gel to solidify. About 80-100,000 ECs were then added to the Matrigel-coated well and cultured for 24 h at 37 °C. Formation of tubular structures on a two-dimensional Matrigel surface was observed after 16 to 24h under an optical microscope.

### CRISPR-Cas9-Mediated Targeting

A) *Pecam1* intron2-enhancer deletion was induced in E14-Tg2a using Alt-R™ CRISPR-Cas9 System (IDT) following the manufacturer’s specifications. This genome editing system is based on the use of a ribonucleoprotein (RNP) consisting of *S. pyogenes* Cas9 nuclease complexed with guide RNA (crRNA:tracrRNA duplex). The crRNA is a 20 nt custom synthesized sequence that is specific for the target and contains a 16 nt sequence that is complementary to the tracrRNA. The specific crRNA sequences were: Pecam1_int2-crRNA1 and Pecam_int2-crRNA3 (sequences shown in Additional file 5: Tab. S5). CRISPR-Cas9 tracrRNA-ATTO 550 (5 nmol catalogue no. 1075927) is a conserved 67 nt RNA sequence that is required for complexing to the crRNA so as to form the guide RNA that is recognized by S.p. Cas9 (Alt-R S.p. Cas9 Nuclease 3NLS, 100 μg catalogue no. 1081058). The fluorescently labelled tracrRNA with ATTO™ 550 fluorescent dye is used to FACS-purify transfected cells. The protocol involves three steps: (1) annealing of the crRNA and tracrRNA, (2) assembly of the Cas9 protein with the annealed crRNA and tracrRNAs, and (3) delivery of the ribonucleoprotein (RNP) complex into mESC by reverse transfection. Briefly, we annealed equimolar amounts of resuspended crRNA and tracrRNA to a final concentration (duplex) of 1 μM by heating at 95°C for 5 min and then cooling to room temperature. The RNA duplexes were then complexed with Alt-R S.p. Cas9 enzyme in OptiMEM media to form the RNP complex, which was then transfected into mESCs using the RNAiMAX transfection reagent (Invitrogen #13778-150). After 48h incubation, cells were trypsinized and ATTO 550 +, transfected cells were purified by FACS.

Fluorescent cells (approximately 65% of the total cell population) were plated at very low density to facilitate colony picking. We picked and screened by PCR 96 clones. Primer sequences are indicated in the Additional file 5: Tab. S5. Positive clones were confirmed by DNA sequencing.

B) For the *Notch1* intron15-enhancer deletion we followed the same procedure but using different target sequences: Notch1_int15-crRNA1 and Notch1_int15-crRNA3 (sequences shown in the Additional file 5: Tab. S5).

### Generation of dCas9-LSD1 expressing mESC line

20 µg of plasmid p-dCas9-LSD1-Hygro (a gift from Stephan Beck and Anna Koeferle, available through Addgene plasmid #104406; http://n2t.net/addgene:104406; RRID:Addgene_104406) was linearized with AhdI enzyme and electroporated in mESC (1×10^7^ cells/10cm plate). The electroporation parameters used were 0.24 kV and 500 μF. The cells were maintained in Hygromicin B selection (500µg/ml) for 10 days. Individual colonies were isolated, expanded and screened by PCR for inserted sequence for both DNA and RNA. Primer sequences are in the Additional file 5: Tab. S5.

### CRISPR-dCas9:LSD1-Mediated epigenetic reprogramming strategy

Epigenetic targeting of putative enhancer elements was induced by transfection of dCas9-LSD1-expressing mESC line with specific gRNA complex (crRNA:tracrRNA duplex). For each enhancer element we designed 3 crRNA sequences (shown in Additional file 5: Tab. S5).

Then, we annealed equimolar amounts of resuspended crRNA and tracrRNA labelled with ATTO™ 550 fluorescent dye to a final concentration (duplex) of 1 μM by heating at 95°C for 5 min and then cooling to room temperature. For gRNAtransfection, cells were plated at 8×10^5^ per well in six-well plates and transfected with gRNA complex (crRNA:tracrRNA 10nM) in antibiotic-free medium using Lipofectamine RNA iMAX Reagent (Invitrogen #13778-150), according to the instructions of the manufacturer. 24 hours after transfection, fluorescent ATTO 550+ cells (approximately 90-95% of the total cell population) were harvested and subjected to the differentiation protocol. crRNA sequences are listed in the Additional file 5: Tab. S5.

### Flow cytometry

We dissociated cells with Trypsin-EDTA or with the Embryoid Body dissociation kit (cod. 130-096-348 Miltenyi Biotec). Dissociated cells (1×10^6^ cells/100 μl) were incubated with primary antibodies (VE-Cadherin-APC, mouse cod.130-102-738) directly conjugated (1:10) in PBS-BE solution (PBS, 0.5% BSA, 5 mM EDTA) for 20 min on ice. Subsequently, cells were washed twice with 2 ml of PBS-BE. Cells were analyzed using the BD FACS ARIAIII™ cell sorter. Negative controls were incubated with fluorochrome-labeled irrelevant isotype control antibody (REA Control APC, mouse cod. 130-113-446 Miltenyi Biotec).

### Quantitative RT-PCR

Total RNA was isolated from mouse ESCs with QIAzol lysis reagent (Qiagen #79306), according to the manufacturer’s protocol. The isolated RNAs were quantified using a NanoDrop spectrophotometer 1000. Before reverse transcription, RNA samples were treated with DNAse I to eliminate any contamination with genomic DNA. cDNA was transcribed using 1 or 2 μg total RNA with the High-Capacity cDNA reverse transcription kit (Applied Biosystem catalog. n. 4368814). cDNAs were amplified using myTaq™ DNA polymerase (Meridian Bioscience) and a standard 3-step cycling PCR profile: 10 min at 94°C, 30 amplification cycles (denaturation at 94°C for 30sec, annealing at 60°C for 30sec, and extension at 72°C for 30sec), followed by a final extension at 72°C for 10 min. Quantitative gene expression analyses (qRT-PCR) were performed using PowerUp™ SYBR Green Master Mix (Applied Biosystem #A25742). Relative gene expression was evaluated using the “2^-ΔCt^” method, and *Gapdh* expression as normalizer. cDNA was amplified by qRT-PCR, using StepOnePlus™ Real-Time PCR System. The run used was holding stage (95°C - 10min); cycling stage (95°C – 15 sec, 60°C – 1min for 40 cycles); melt curve stage (95°C – 15sec, 60°C – 1min, 95°C – 15sec). The cycle threshold (Ct) was determined during geometric phase of the PCR amplification plots, as illustrated in the manufacturer’s protocol. Expression data are shown as the mean ± SD. Primer sequences are listed in Supplementary Table 5. Graph Pad Prism software v8.00 (GraphPad) was used to analysis of qRT-PCR data. Relative mRNA levels were analysed in triplicate and data were presented as means ± SD. Two-way repeated measures ANOVA test (ANOVA 2way-RM) was used to assess statistically significant interaction effect between “time” and “genotype” on a gene expression variable. Other two statistical methods between groups of data were used: nonparametric and parametric test. The first was nonparametric Wilcoxon matched-pairs signed rank test, one-tailed; the second statistical analysis were performed using the parametric Student’s paired t-test, one-tailed. Shapiro-Wilk test was performed to determine the normality distribution of the dataset.

### RNA-seq

Total RNA was isolated from d2 (n. 2 biological replicates) and d4 (n. 2 biological replicates) cells with QIAzol lysis reagent (Qiagen #79306), according to the manufacturer’s protocol. RNA concentration was estimated using a Nanodrop spectrophotometer 1000. Libraries were prepared according to the Illumina strand specific RNA-seq protocol. Libraries were sequenced on the Illumina platform NextSeq 500, in paired end, 75 bp reads.

### ATAC-seq

mESCs were collected at day2 and day4 and then washed two times in PBS, harvested, counted using a hemacytometer chamber and pelleted. 15000 cells/sample for mESC were treated with Tagment DNA Buffer 2x reaction buffer with Tagment DNA Enzyme (Illumina) according to the manufacturer’s protocol. After washes in PBS, cells were suspended in 50 mL of cold lysis buffer (10 mM Tris-HCl, pH 7.4, 10 mM NaCl, 3 mM MgCl2, 0.1% IGEPAL CA-630) and immediately spun down at 500 × g for 10 min at 4°C. Fresh nuclei were treated with Transposition mix and Purification (Illumina #FC121-130), the nuclei were incubated at 37°C in Transposition Reaction Mix (25 µL reaction buffer, 2.5 µL Transposase, 22.5 µL Nuclease free water), purified using Qiagen MinElute PCR Purification Kit (catalog no./ID: 28006) and eluted in 10 µL of nuclease free water. Sequencing library was prepared from the fragmented amplified tagmented DNA. Fragmentation size was evaluated using the Agilent 4200 TapeStation. Two biological replicates for each condition were sequenced using Illumina NextSeq500 system to obtain paired-end (PE) reads of 60 bp.

### Sequence data analysis

For RNA-seq sequencing data, we assessed the quality of the paired-end (PE) reads of length 75bp using FastQC. We filtered the low-quality and short PE reads and trimmed the universal Illumina adapters using cutadapt (v2.9) [28] by setting the following parameters: -q 30 -m 30. Post-trimming, we re-assessed the quality and compiled the report using multiQC. We aligned the PE reads to mm10/GRCm38 reference genome (primary assembly) using STAR aligner (v2.6.0a) [29] following two steps: i) Generation of Ensembl mm10 reference genome index (release 102) setting the parameters --sjdbGTFfile 100 --sjdbOverhang 100 ii) Alignment of PE reads to the reference genome (--sjdbOverhang 100 --quantMode GeneCounts --outSAMtype BAM SortedByCoordinate). We provided the sorted BAM as input and the mm10 (GRCm38.p4) primary assembly annotation file in GTF format(https://www.gencodegenes.org/mouse/release_M10.html) as reference annotation to quantify the gene expression levels with the featureCounts function from the Rsubread package (v2.0.1) [30] (annot.ext=”mm10.v102’’ gtf file, useMetaFeatures=TRUE, allowMultiOverlap=FALSE, strandSpecific=2, CountMultiMappingReads=FALSE). After that, we retained the expressed gene matrix by filtering out from the read count matrix the zero and low count genes (CPM < 0.5) using the proportion test method from the NOIseq package (v2.34.0) [31]. Then, we processed the expressed gene matrix using NOIseq (v2.34.0), obtaining a set of normalized read counts (Upper Quartile (UQUA) and identified differentially expressed genes (DE) using the noiseq function and setting the posterior probability value cutoff > 0.8. Simultaneously, we normalized the expressed count matrix with the sizefactor function in DEseq2 [32] and we used the DESeq function with default options, then we selected the DE genes by setting the adj.p-value cutoff < 0.01. Finally, we considered DE genes in our study, those genes that were declared DEs from both methods. We performed gene onthology enrichment analysis using the gProfiler2 R package (v0.2.2) [33], providing the common DE gene list as input, the expressed gene list as background, and setting Benjamini-Hochberg FDR (BH-FDR) cutoff to < 0.01.

ATAC-seq sequences underwent quality control (using FastQC and multiQC), adapter trimming, and filtering using cutadapt (v2.9) with parameters -q 30 --m 30 and universal Illumina adapters. Then, we aligned the PE sequences to the mouse genome (mm10) containing only canonical chromosomes with Bowtie2 (v2.3.4.3) [34], setting the options -q -t --end-to-end --very-sensitive -X 1000. After removing reads mapping to the mitochondrial chromosome, we removed duplicates and reads mapping to multiple positions using sambamba (v0.6.8) with -F “[XS] == null and not unmapped and not duplicate” [35]. We called the ATAC peaks for each sample using MACS2 (v2.1) [36] with the option -BAMPE –nomodel –shif100 –extsize 200, which are the suggested parameters to handle the Tn5 transposase cut site. After that, we removed those peaks overlapping the mm10 blacklist regions (downloaded from https://github.com/Boyle-Lab/Blacklist/blob/master/lists/mm10-blacklist.v2.bed.gz) using samtools [37]. Then, for each condition, we defined the consensus lists of enriched regions as the peak regions common to both replicates using the intersectBed function from the BedTools v2.26.0 [38]. To identify differentially enriched regions (DARs) between d4 and d2, we selected the regions consistently enriched or decreased in DEScan2 (v1.18.2) [39] and confirmed with the same sign also in DiffBind (v3.8.4) [40]. For DEScan2, we first created a peak-set using the finalRegions function (zThreshold = 1, minCarriers = 2) after loading all of the MACS2 peaks not overlapping black list regions. Then, we called the DARS with the edgeR [41] method as follows: we estimated the count dispersion using the estimateDisp function, we fitted the robust glmQLFit model and used the glmQLFTest, with the default parameters, setting the adj. p-value to 0.01. For DiffBind (v3.8.4), we set dba.count(minOverlap = 2), dba.contrast(minMembers = 2), dba.analyze(method = DBA_EDGER) and 0.01 as adj. p-value. Finally, we selected the DARs regions of DEScan2 confirmed by DiffBind using subsetByOverlaps in GenomicRanges [42]. We annotated the consensus peak lists, the common peak set, and DARs to genes using the makeTxDbFromEnsembl function from ChIPseeker (v1.29.1) [43] by associating to each peak/region the nearest gene, setting the TSS region [−1000, 1000] and using them the release 102 from the *mus musculus* Ensemble database.

Finally, we selected the chromatin-enriched regions at d4 with annotated nearest genes intersecting some lists of marker genes from the single cell experiment described in [20], i) the marker genes of endothelial cell clusters in *Tbx1^Cre^* Ctrl and cKO embryos at E9.5 (Supplementary Data 5, cluster 6 of the cited publication); ii) the marker genes of endothelial cell clusters in *Mesp1^Cre^* Ctrl and cKO at E9.5 (Supplementary Data 3, cluster 2); iii) Marker genes of endothelial cell clusters in *Mesp1^Cre^* Ctrl and cKO at E9.5 (Supplementary Data 3, cluster 16). We used such regions to identify enriched motifs using HOMER (Hypergeometric Optimization of Motif EnRichment) [44]. From the motif list of regions associated with genes intersecting marker genes of endothelial cell clusters in *Tbx1^Cre^*, we selected GATA3 and ERG motifs, we identified those regions containing both motifs and we calculated the percentage. We performed the enhancer prediction of regions associated with genes intersecting the list of marker genes of endothelial cell cluster 6 in *Tbx1^Cre^*Ctrl embryos at E9.5 as described in the next section. Finally, we merged the list of regions derived from the intersections of d4 DARs with annotated nearest genes intersecting lists of marker genes of Mesp1cre c2 and c16 clusters and selected unique regions, then we performed the enhancer prediction using these regions.

### Enhancer prediction

We implemented a machine learning approach to assign a probability score of being enhancers to peak regions from ATAC-seq data using the logistic regression model with the L-1 penalty. We performed all analyses using Rstudio and R version 4.2.0 (https://www.r-project.org/).

First, we downloaded the coordinates (chromosome, start-end position) of the 695 enhancers marked as positive from the VISTA ENHANCER Browser (http://enhancer.lbl.gov). Then, since the enhancers’ coordinates were in mm9, we mapped them into mm10 using the lift-over function (https://genome.ucsc.edu/cgi-bin/hgLiftOver). Next, we created 695 non-enhancer regions to use as negative examples. For this purpose, we randomly shuffled the genome to get genomic coordinates that do not overlap the positive enhancer coordinates with the shuffle function from the BedTools v2.26.0 [38] using the parameters -g mm10 -excl the positive enhancer file merged with mm10 blacklist regions file. (https://github.com/Boyle-Lab/Blacklist/blob/master/lists/mm10-blacklist.v2.bed.gz). Such non-enhancer regions have the same lengths as the positive enhancers. Finally, we built a binary response vector with 1390 components assigning 1 to the positive enhancers and 0 to the negative enhancers. Second, we downloaded 385 datasets using Chipseeker (v1.36) R package with the function ChIPseeker::downloadGEObedFiles(genome= “mm10”) (https://bioconductor.org/packages/release/bioc/html/ChIPseeker.html) containing peak coordinates of histone modifications, p300 and CTCF transcription factors, at different cell states and cell types (in mm10). After merging the replicates using intersectBed function from the BedTools, we got 327 files. Then, we filtered out the datasets containing KO experiments or other treatments. Overall, we obtained 81 epigenetic tracks of peaks in bed format and 2 in bedgraph format. Next, we intersected the 1390 enhancer and non-enhancer regions with the 81 epigenetic tracks in bed format using the Findoverlaps function from the GenomicRanges R package (v1.52.0) (https://bioconductor.org/packages/release/bioc/html/GenomicRanges.html). Finally, we built a binary matrix of dimension 1390×81 where we assigned 1 at the positions with an overlap and 0 where they do not. For the remaining 2 epigenetics tracks in bedgraph format, we used the sum of the coverage at each of the 1390 regions. The feature matrix of dimension 1390×83 is formed by combining the two parts. We split the dataset into the Training-set (80% of the 1390 regions) and the Test set (20% of the 1390 regions). Then, we trained the L1-penalized Logistic Regression (i.e., Lasso logistic regression) on the training set using K-fold cross-validation to choose the best regularization parameter. To this purpose, we used the glmnet package (v4.1.7) (https://cran.r-project.org/web/packages/glmnet/index.html) with the command: cv.out=cv.glmnet(X.train, Y.train,alpha=1, family= “binomial”). After that, we fitted the training set using the lasso.mod.train= glmnet(X.train, Y.train, lambda = bestlam, alpha = 1, family =“binomial”), where bestlam is the regularization parameter obtained from the cross-validation. In the validation phase, we used the assess.glmnet() function to determine the accuracy of the test-set. Finally, we predicted the scores (i.e., the probability of being an enhancer) to the genomic coordinates of interest. Since the scores vary between 0 (corresponding to minimum probability) and 1 (corresponding to maximum similarity), we used the threshold of 0.5 to establish if a given region is an enhancer. The whole procedure, starting from the random generation of the 695 non-enhancer regions to the final prediction, was repeated 10 times, and the final prediction consisted of the average of the individual prediction scores over which we applied the threshold of 0.5.

## DECLARATIONS

### Ethics approval and consent to participate

N/A

### Consent for publication

N/A

### Availability of data and materials

Data supporting the results are included in the figures, additional files, and in the GEO database under the accession number GSE235651 (RNA-seq and ATAC-seq data).

### Competing interests

The authors declare that they have no competing interests.

## Funding

This work was funded by grants from the Fondation Leducq 15CVD01 (AB and EI) and the Italian Ministry of University and Research PRIN 20179J2P9J (AB and EI).

## Authors’ contributions

IA: performed experiments, assembled figures, provided conceptual input to experimental design and edited the manuscript; OL and VPK: performed bioinformatic analysis, assembled figures, and contributed to manuscript editing; AC: performed experiments; SA: performed experiments; GL: performed experiments, provided experimental design input and contributed to manuscript editing; RF: performed experiments; CA: supervised bioinformatic analysis, provided conceptual input, and contributed to manuscript editing; EI: provided funding, contributed conceptual input to experimental design, and edited the manuscript; AB: provided funding, designed the experimental plan, wrote the manuscript.

## Acknowledgements

We thank Marchesa Bilio for technical help; Laura Pisapia and Enzo Mercadante at the flow cytometry core of the Institute of Genetics and Biophysics. RNA-seq and ATAC-seq samples were sequenced by Genomix4Life SrL, Salerno, Italy.

## ADDITIONAL FILES

### Additional file 1: Table S1

Sheet 1: Differentially expressed genes (d4 Vs d2). Positive values are for genes up regulated at d4.

Sheet 2: All expressed genes at d2 and at d4.

Sheet 3: GO_RNAseq_DEGs.

Sheet 4: GO_RNAseq_upreg at d4.

### Additional file 2: Table S2

Sheet 1: Differentially accessible regions (DARs), d4 Vs d2. Positive values are for regions more accessible at d4.

Sheet 2: All consensus ATAC peaks at d2.

Sheet 3: All consensus ATAC peaks at d4.

Sheet 4: Common peak set (peakome).

### Additional file 3: Table S3

Sheet 1: 252 marker genes from *Tbx1^Cre^* EC cluster 6.

Sheet 2: 1217 marker genes from *Mesp1^Cre^* cluster c2.

Sheet 3: 1434 marker genes from *Mesp1^Cre^* cluster c16.

Sheet 4: 536 unique DARs mapped to EC marker genes from *Mesp1^Cre^* clusters.

### Additional file 4: Table S4

Sheet 1: Motifs *Tbx1^Cre^* EC cluster 6.

Sheet 2: Motifs *Mesp1^Cre^* EC cluster c2.

Sheet 3: Motifs *Mesp1^Cre^* EC cluster c16.

### Additional file 5: Table S5

Primers and gRNA sequences

## Notes

### Competing Interest Statement

The authors have declared no competing interest.

